# Laminar architecture of visual responses in supplementary eye field of macaques

**DOI:** 10.1101/2023.10.03.560770

**Authors:** Pranavan Thirunavukkarasu, Steven P. Errington, Amirsaman Sajad, Jeffrey D. Schall

## Abstract

Previously, we have described the laminar organization of neurons in the supplementary eye field (SEF) that signal error, reward gain and loss, conflict, event timing, and goal maintenance. Here we describe the laminar organization of visually responsive neurons that were active during performance of a saccade stop-signal task. Nearly 40% of isolated neurons exhibited enhanced or suppressed responses to a visual target for a potential saccade, with the majority exhibiting enhanced activity and three-quarters with broad spikes. Visually responsive neurons were observed in all layers but were less common in layers 5 and 6. Response latencies were comparable to those reported previously, which are significantly later than those measured in occipital and temporal visual areas but overlapping those measured in cingulate cortex. Task-related visual response latency varied across cortical layers. Response latency was significantly earlier for neurons with narrow spikes. Neurons with task-related visual responses discharged until after saccade production. Around three-fifths of visually responsive neurons were most sensitive to the visual target appearing in one hemifield. Many neurons in layer 2 had ipsilateral receptive fields. Laminar current-source density aligned on visual target presentation revealed the earliest sink in layers 3 followed by a prolonged strong sink more superficially coupled with a weaker prolonged sink in layer 5 and a transient sink in layer 6. The current sink in layers 2 and 3 was stronger for ipsilateral stimuli. These findings reveal new details about visual processing in medial frontal cortex and complete the first catalogue of laminar organization of functional signals in a frontal lobe area.

## INTRODUCTION

This report continues a series of investigations of the laminar organization of neurons present in SEF— an agranular area on the dorsomedial convexity in macaques heavily involved with visually guided behavior. Previously, we described the laminar organization of passive visual responses (Godlove et al., 2014); here, we describe responses during an active task. We also characterized substantial differences in the laminar functional organization of agranular SEF relative to granular V1 (Ninomiya et al., 2015); here, we describe additional distinguishing features of SEF relative to visual cortical areas. Finally, we have described the laminar organization in SEF of neurons signaling error, reward gain and loss, conflict, event timing, and goal maintenance during a saccade countermanding task (Sajad et al., 2019; Sajad et al., 2022). Here, we describe the laminar profile of task-related visually responsive neurons. Knowing how SEF processes visual information is necessary to understand how environmental attributes are integrated into cognitive control.

## METHODS

### Experimental models and subject details

#### Non-human primates

Data was collected from one male bonnet macaque (Eu, *Macaca Radiata*, 8.8kg) and one female rhesus macaque (X, *Macaca Mulatta*, 6.0 kg) performing a countermanding task (Hanes and Schall, 1995; Godlove et al., 2014). All procedures were approved by the Vanderbilt Institutional Animal Care and Use Committee in accordance with the United States Department of Agriculture and Public Health Service Policy on Humane Care and Use of Laboratory Animals.

### Method details

#### Stop-signal task

The saccade stop-signal (countermanding) task utilized in this study has been widely used previously (Hanes and Schall, 1995; Hanes and Carpenter, 1999; Cabel et al., 2000; Colonius et al., 2001; Kornylo et al., 2003; Morein-Zamir and Kingstone, 2006; Walton and Gandhi, 2006; Thakkar et al., 2011; Thakkar et al., 2015; Godlove and Schall, 2016; Wattiez et al., 2016; Verbruggen et al., 2019). Briefly, trials were initiated when monkeys fixated at a central point. Following a variable time-period, the center of the fixation point was removed leaving an outline. At this point, a peripheral target was presented simultaneously on either the left or right hand of the screen. In this study, one target location was associated with a larger magnitude of fluid reward. The lower magnitude reward ranged from 0 to 50% of the higher magnitude reward amount. This proportion was adjusted to encourage the monkey to continue responding to both targets. The stimulus-response mapping of location-to-high reward changed across blocks of trials. Block length was adjusted to maintain performance at both targets, with the number of trials in each block determined by the number of correct trials performed. In most sessions, the block length was set at 10 to 30 correct trials. Erroneous responses led to repetitions of a target location, ensuring that monkeys did not neglect low-reward targets in favor of high-reward targets – a phenomenon demonstrated in previous implementations of asymmetrically rewarded tasks (Kawagoe et al., 1998).

On most of the trials, the monkey was required to make an eye movement to this target (no-stop trials). However, on a proportion of trials the center of the fixation point was re-illuminated (stop-signal trials); this stop-signal appeared at a variable time after the target had appeared (stop-signal delay; SSDs). An initial set of SSDs, separated by either 40 or 60 ms, were selected for each recording session. The delay was then manipulated through an adaptive staircasing procedure in which stopping difficulty was based on performance. When a subject failed to inhibit a response, the SSD was decreased by a random step to increase the likelihood of success on the next stop trial.

Similarly, when subjects were successful in their inhibition, the SSD was increased to reduce the likelihood of success on the next stop trial. This procedure was employed to ensure that subjects failed to inhibit action on approximately 50% of all stop-signal trials. On no-stop trials, the monkey was rewarded for making a saccade to the target. On stop-signal trials, the monkey was rewarded for withholding the saccade and maintaining fixation on the fixation spot. Following a correct response, an auditory tone was sounded 600ms later, and followed by a high or low fluid reward, depending on the stimulus-response mapping.

#### Animal care and surgical procedures

Surgical details have been described previously (Godlove et al., 2011). Briefly, magnetic resonance images (MRIs) were acquired with a Philips Intera Achieva 3T scanner using SENSE Flex-S surface coils placed above or below the animal’s head. T1-weighted gradient-echo structural images were obtained with a 3D turbo field echo anatomical sequence (TR = 8.729ms; 130 slices, 0.70mm thickness). These images were used to ensure Cilux recording chambers were placed in the correct area (Crist Instruments). Chambers were implanted normal to the cortex (Monkey Eu: 17°; Monkey X: 9°; relative to stereotaxic vertical) centered on midline, 30mm (Monkey Eu) and 28mm (Monkey X) anterior to the interaural line.

#### Cortical mapping and electrode placement

Chambers implanted over the medial frontal cortex were mapped using tungsten microelectrodes (2-4 MΩ, FHC, Bowdoin, ME) to apply 200ms trains of biphasic micro-stimulation (333 Hz, 200 µs pulse width). The SEF was identified as the area from which saccades could be elicited using < 50 µA of current (Schall, 1991b; Thompson et al., 1996; Martinez-Trujillo et al., 2004). In both monkeys, the SEF chamber was placed over the left hemisphere.

A total of five penetrations were made into the cortex: two in monkey Eu, and three in monkey X. Three of these penetrations were perpendicular to the cortex. In monkey Eu, the perpendicular penetrations sampled activity at site P1, located 5mm lateral to the midline and 31mm anterior to the interaural line. In monkey X, the perpendicular penetrations sampled activity at site P2 and P3, located 5mm lateral to the midline and 29 and 30mm anterior to the interaural line, respectively. However, during the mapping of the bank of the cortical medial wall, we noted both monkeys had chambers place ∼1mm to the right respective to the midline of the brain. This was confirmed through co-registered CT/MRI data. Subsequently, the stereotaxic estimate placed the electrodes at 4mm lateral to the cortical midline opposed to the skull-based stereotaxic midline.

#### Data acquisition

Spiking activity and local field potentials were recorded from five sites within the SEF using a 24-channel Plexon U-probe (Dallas, TX) with 150 µm interelectrode spacing allowing sampling from all layers. Penetrations in three of these sites were perpendicular to the cortex. The U-probes were 100 mm in length with 30 mm reinforced tubing, 210 µm probe diameter, 30° tip angle, and had 500 µm between the tip and first contact. Contacts were referenced to the probe shaft and grounded to the headpost. We used custom built guide tubes consisting of 26-gauge polyether ether ketone (PEEK) tubing (Plastics One, Roanoke, VA) cut to length and glued into 19-gauge stainless steel hypodermic tubing (Small Parts Inc., Logansport, IN). This tubing had been cut to length, deburred, and polished so that they effectively support the U-probes as they penetrated dura and entered cortex. The stainless-steel guide tube provided mechanical support, while the PEEK tubing electrically insulated the shaft of the U-probe, and provided an inert, low-friction interface that aided in loading and penetration.

Microdrive adapters were fit to recording chambers with < 400 µm of tolerance and locked in place at a single radial orientation (Crist Instruments, Hagerstown, MD). After setting up hydraulic microdrives (FHC, Bowdoin, ME) on these adapters, pivot points were locked in place by means of a custom mechanical clamp. Neither guide tubes nor U-probes were removed from the microdrives once recording commenced within a single monkey. These methods ensured that we were able to sample neural activity from precisely the same location relative to the chamber on repeated sessions.

Electrophysiology data were processed with unity-gain high-input impedance head stages (HST/32o25-36P-TR, Plexon). All data were streamed to a single data acquisition system (MAP, Plexon, Dallas, TX). Time stamps of trial events were recorded at 500 Hz. Eye position data were streamed to the Plexon computer at 1 kHz using an EyeLink 1000 infrared eye-tracking system (SR Research, Kanata, Ontario, Canada).

#### Cortical depth assignment

The retrospective depth of the electrode array relative to grey matter was assessed through the alignment of several physiological measures. Firstly, the pulse artifact was observed on a superficial channel which indicated where the electrode was in contact with either the dura mater or epidural saline in the recording chamber; these pulsated visibly in synchronization with the heartbeat. Secondly, a marked increase of power in the gamma frequency range (40-80Hz) was observed at several electrode contacts, across all sessions. Previous literature has demonstrated elevated gamma power in superficial and middle layers relative to deeper layers (Maier et al., 2010; Xing et al., 2012; Smith and Sommer, 2013). Thirdly, an automated depth alignment procedure was employed which maximized the similarity of CSD profiles evoked by passive visual stimulation between sessions (Godlove et al., 2014).

#### Data collection protocol

An identical daily recording protocol across monkeys and sessions was carried out. In each session, the monkey sat in an enclosed primate chair with their head restrained 45 cm from a CRT monitor (Dell P1130, background luminance of 0.10 cd/m2). The monitor had a refresh rate of 70 Hz, and the screen subtended 46° x 36° of the visual angle. Eye position data was collected at 1 kHz using an EyeLink 1000 infrared eye-tracking system (SR Research, Kanata, Ontario, Canada). This was streamed to a single data acquisition system (MAP, Plexon, Dallas, TX) and amalgamated with other behavioral and neurophysiological data. After advancing the electrode array to the desired depth, they were left for 3 to 4 hours until recordings stabilized across contacts. This led to consistently stable recordings. Once these recordings stabilized, an hour of resting-state activity in near-total darkness was recorded. This was followed by the passive presentation of visual flashes followed by periods of total darkness in alternating blocks. Finally, the monkey then performed approximately 2000 to 3000 trials of the saccade countermanding (stop-signal) task.

#### Data Analysis

Significant modulation was defined as activity that was 3 SD greater than a baseline period (−250 to 0 ms relative to target) for longer than 75 ms and reached a magnitude of 6 SD for at least 25 ms. The visual period was 50 to 200 ms, post-target. All statistical procedures for neural data analysis were done using two-tailed tests unless otherwise specified. All statistics were performed using MATLAB 2022b/2023a (MathWorks Inc; Natick, MA, USA)

#### Current Source Density

We used the same methods reported in Herrera et al. (2022) to calculate the CSD. We computed the CSD signal using the spline-iCSD method (Pettersen et al., 2006) as implemented in the CSDplotter toolbox (https://github.com/espenhgn/CSDplotter) with custom MATLAB scripts. For the standard CSD method, CSD was computed from the raw signal by taking the second spatial derivative along electrodes (Nicholson and Freeman, 1975; Schroeder et al., 1998; Mehta et al., 2000; Westerberg et al., 2019) and converting voltage to current based on estimates of impedance (Logothetis et al., 2007). We computed the CSD by taking the second spatial derivative of the LFP:

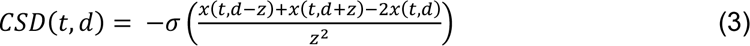

where *x* is the extracellular voltage at time *t* measured at an electrode contact at depth *d*, z is the inter-electrode distance, and σ is conductivity. CSD was baseline corrected at the trial level by subtracting the average activation during the 300 ms preceding array onset from the response at all timepoints. CSD was clipped 10 ms before saccade at the trial level to eliminate the influence of eye movements.

## RESULTS

We acquired 33,816 trials across 29 sessions from two macaques (Eu: 11,583 trials; X: 22,233 trials) performing the saccade countermanding task (**Fig. 1A**). Both monkeys exhibited typical countermanding behavior, details of which have been previously reported (Sajad et al., 2019, 2022). Using linear microelectrode arrays (**Fig. 1B**), we isolated 575 neurons (Eu: 331, X: 244) from these sessions.

**Figure 1.**
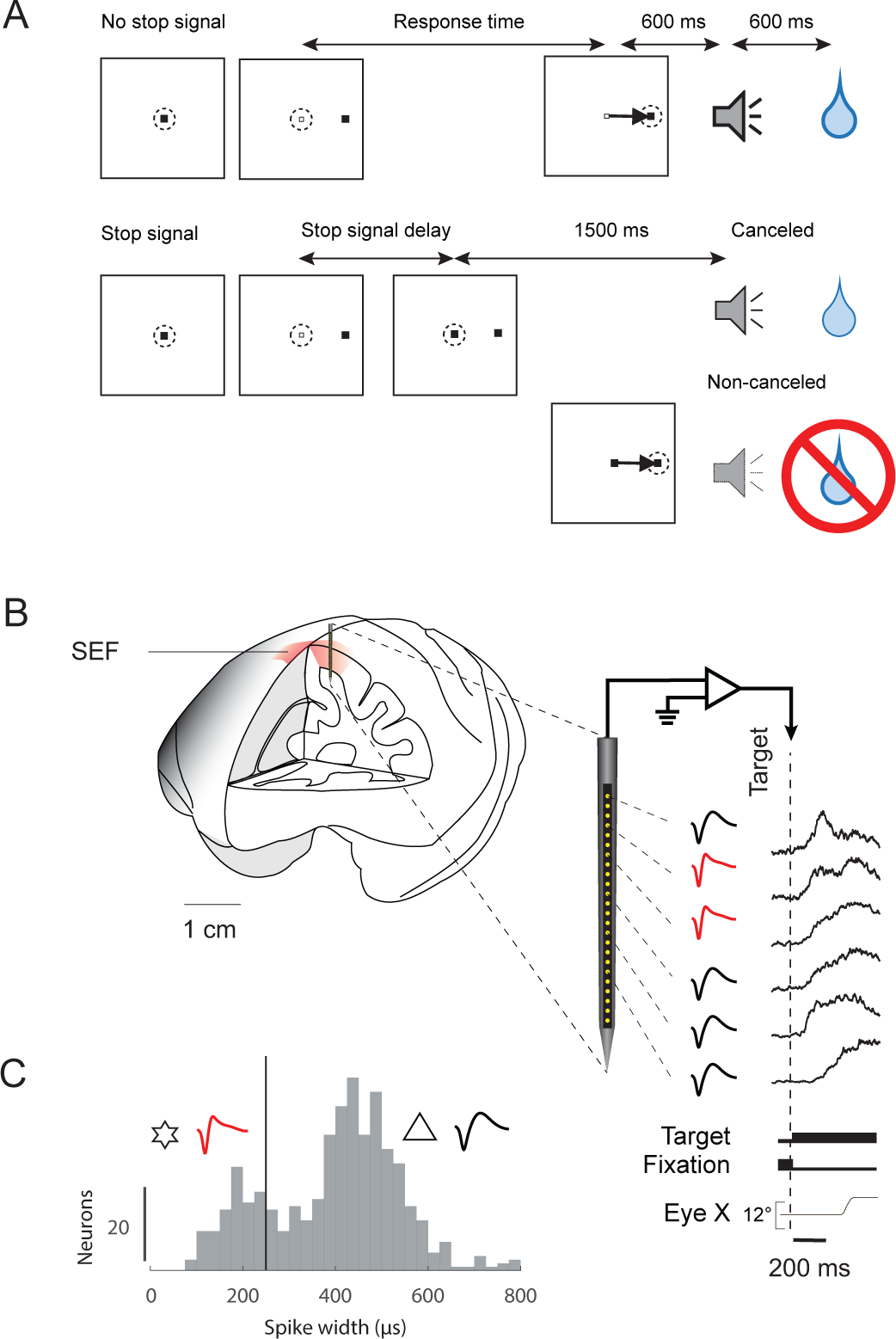
A. Visually-guided saccade-countermanding task. Monkeys initiated trials by fixating on a central point. After a variable time, the center of the fixation point was extinguished. A visual target was presented simultaneously at one of two possible locations. On no-stop-signal trials, monkeys were required to shift gaze to the target, whereupon after 600 ± 0 ms, a high-pitched auditory feedback tone was delivered, and 600 ± 0 ms later, fluid reward was provided. On stop signal trials (∼40% of trials), after the target appeared, the center of the fixation point was re-illuminated after a variable stop-signal delay, which instructed the monkey to cancel the saccade, in which case the same high-pitched tone was presented after a 1,500 ± 0 ms hold time followed, after 600 ± 0 ms, by fluid reward. Stop-signal delay was adjusted such that monkeys successfully canceled the saccade in ∼50% of trials. In the remaining trials, monkeys made noncanceled errors, which were followed after 600 ± 0 ms by a low-pitched tone, and no reward was delivered. Monkeys could not initiate trials earlier after errors. B. Neural spiking was recorded across all layers of SEF using a Plexon U-probe. Neurons with both broad (black) and narrow (red) spikes were sampled. Spiking modulation was measured relative to presentation of the visual target. C. Distributions of spike widths of all neurons sampled (n = 575). The vertical line marks the 250 μs separation criterion used.

### Functional properties of visual responses in supplementary eye field

To identify visually responsive neurons within our sample, we examined the activity around the time the fixation point was extinguished, and the peripheral target, which was the go-signal, was presented. We focused on no-stop trials, in which monkeys were simply required to generate a saccade to the presented target. As summarized in **Table 1**, we observed 226 (∼39%) neurons demonstrated significant deviation from baseline following the presentation of a visual target. Of these, 203 (∼90%) produced a significant increase in firing rates, whilst 23 (∼10%) had a significant decrease in firing rates (**Fig. 2**).

**Figure 2.**
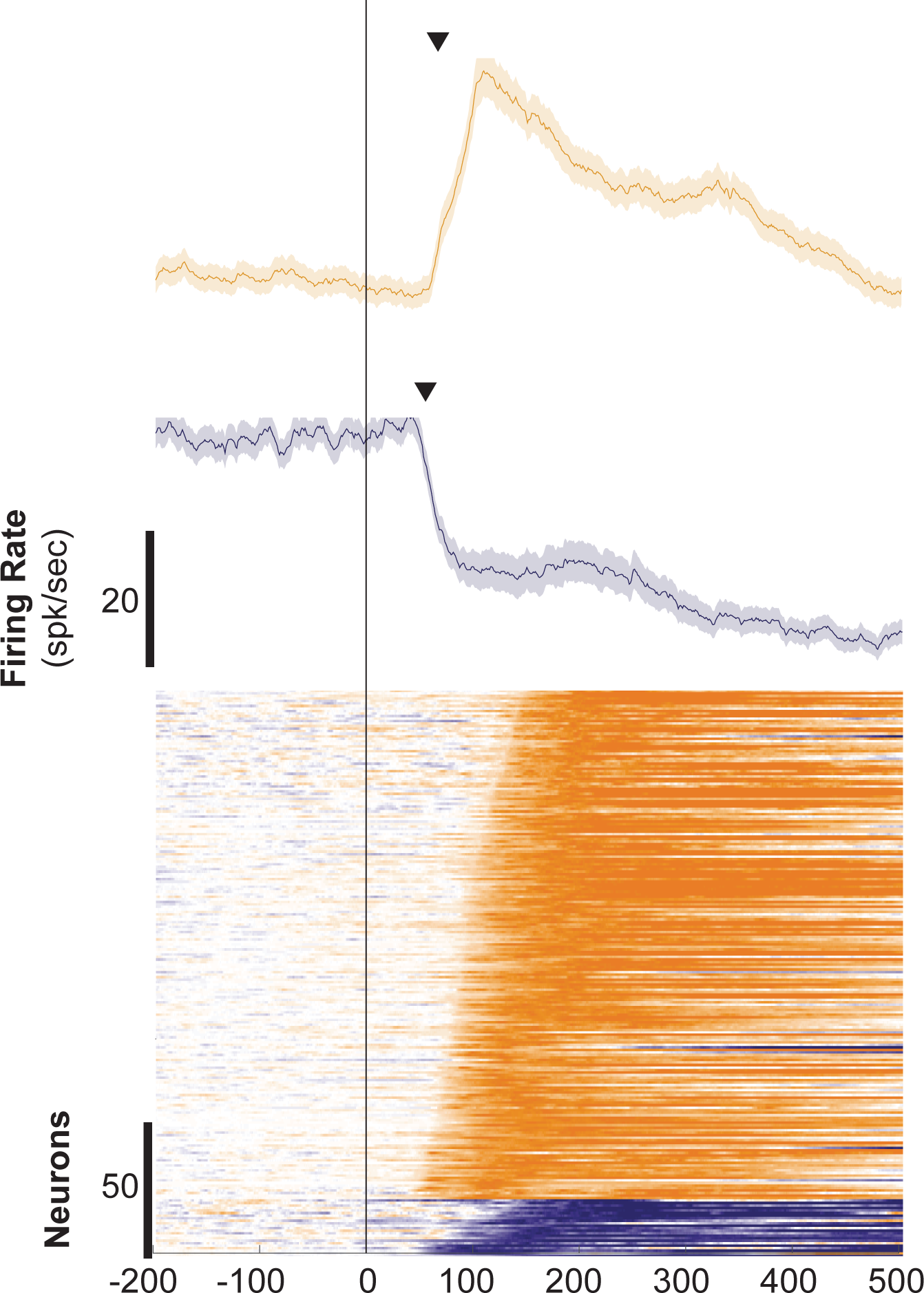
A. Raster and average spike density of representative neurons with facilitated (top) and suppressed (bottom) visual responses. Measured visual latency indicated by arrowhead. B. Color-coded raster of all facilitated (orange) and suppressed (blue) visually responsive neurons ordered by visual latency. C. Proportions of neurons with facilitated, suppressed, or no visual response with broad and narrow spikes.

**Table 1.** Count of neurons recorded from SEF organized by functional properties and separated by spike width.

| Cell Type | *n*<sub>Broad</sub> | *n*<sub>narrow</sub> | *N*<sub>Total</sub><br>(% of Total) |
| --- | --- | --- | --- |
| Non-visual | 287 | 62 | 349<br>(60.7%) |
| Visual |  |  | 226<br>(39.3%) |
| Suppressed | 17 | 6 | 23<br>(4%) |
| Facilitated | 159 | 44 | 203<br>(35.3%) |
| Visual Field Responsive |  |  | 121<br>(21.0%) |
| Ipsi | 56 | 17 | 73<br>(12.7%) |
| Contra | 45 | 3 | 48<br>(8.3%) |

We distinguished neurons with narrow and broad spikes. Spike width was defined as the trough to peak interval of the waveform. The distribution of spike widths within this sample was bimodal (calibrated Hartigan’s dip test, p < 0.01) (**Fig. 1C**). In line with other studies, we employed a conservative cut-off of 250 μs in differentiating cells with narrow or broad spikes (Cohen et al., 2009; Thiele et al., 2016). Using this criterion, 112 units had narrow spikes, and 463 units, broad. This is consistent with a general 1:4 ratio of narrow- and broad-spiking cortical neurons (Dzaja et al., 2014; Ardid et al., 2015). Putatively, narrow-spiking neurons were associated with inhibitory interneurons and broad-spiking neurons with pyramidal neurons (Lemon et al., 2021). Among facilitated neurons, 159 (78%) had broad spikes and 44 (22%), narrow spikes. Among suppressed neurons, 17 (74%) had broad spikes and 6 (26%) had narrow spikes (**Table 1**). Given the few suppressed neurons observed, we will focus on the properties of facilitated neurons.

The visually-evoked activity terminated following the execution of the saccade (**Fig. 3A**). We quantified the balance of visually-evoked and saccade-related activity by measuring discharge rates in the intervals 50 to 200 ms after visual stimulus presentation and ±50 ms relative to saccade initiation and calculating a visuomotor index (VMI) for each neuron, which was the contrast ratio between visual and saccadic activity:

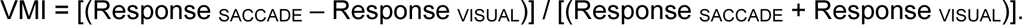

**Figure 3.**
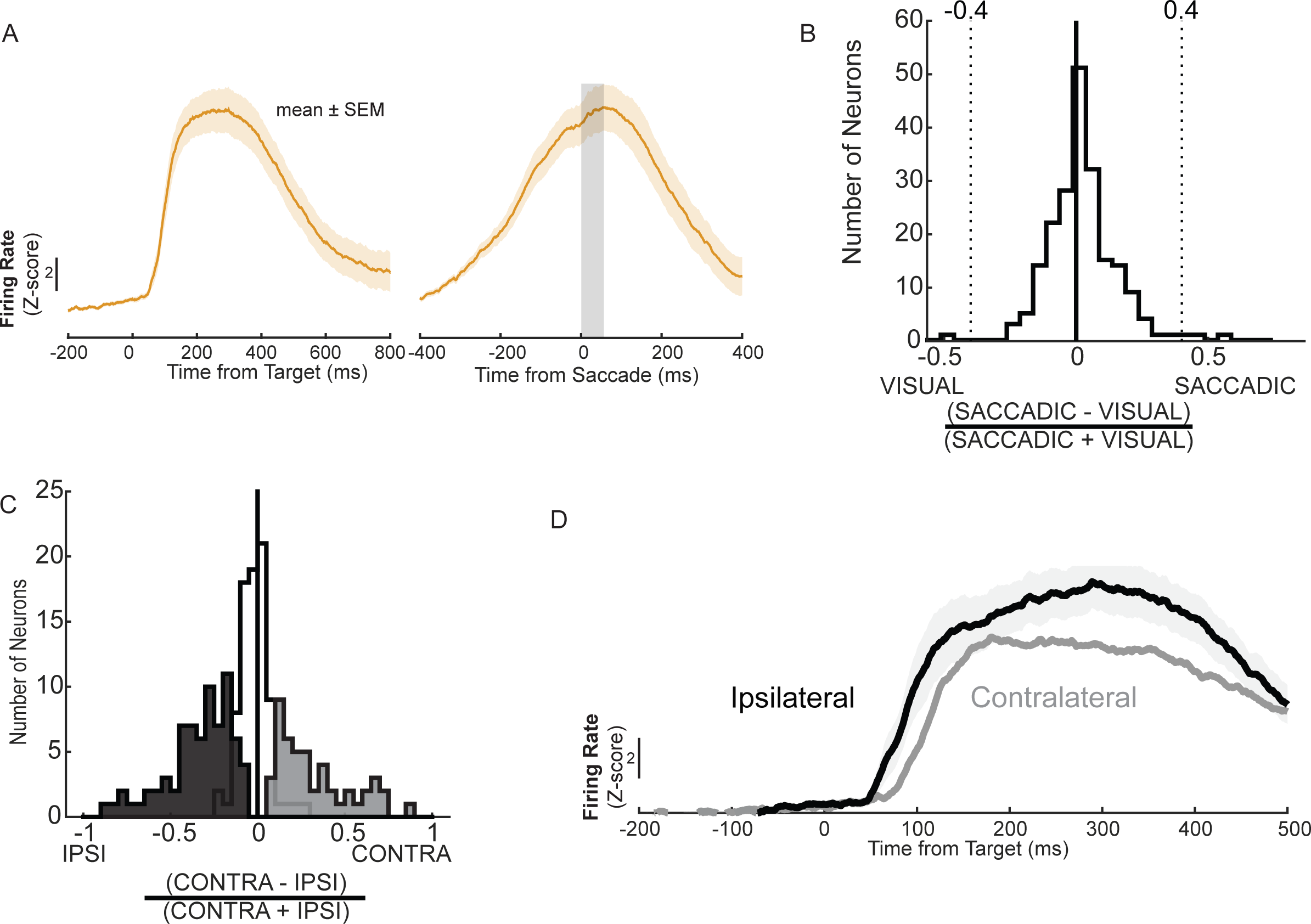
A. Average spike density function of visually responsive neurons aligned on target presentation (left) and saccade initiation (right). B. Histogram of visuomotor index of all neurons with greater saccade-related activity positive and stronger visually evoked activity, negative. Empty bars indicating neurons with equivalent activity and dark bars highlight significant differences where the left dashed black lines correspond to a VMI of −0.4 and the right dashed black line corresponds to a VMI of 0.4. C. Histogram of laterality index with contralateral preferences positive and ipsilateral, negative. Empty bars correspond to neurons displaying equivalent responses and filled bars highlight significant differences. D. Average spike density function of visually responsive neurons for targets in contralateral (light) and ipsilateral (dark) visual fields.

Visuomotor index values span the interval from −1 to +1, with −1 representing greater activity during the visual period and +1, greater activity during the saccadic period. Typically, neurons with VMI between <−0.4 are categorized as visual neurons, VMI > 0.4, as saccadic, and the remainder as visuomotor neurons (Bruce and Goldberg, 1985; Lawrence et al., 2005). All of the visually responsive neurons in SEF were categorized as visuomotor (**Fig 3B**). The distribution of VMI values was significantly greater than 0 (mean VMI = 0.085, t (202) = 9.616, p < 0.001).

We next examined how the visual responses varied with the laterality of the stimuli. Targets appeared at 12° eccentricity on the horizontal meridian in either the contra-or ipsilateral visual field. We quantified the laterality of visually evoked activity by measuring discharge rates in the interval 100 ms from visual facilitation onset and calculating a visual laterality index (VLI) for each neuron, which was the contrast ratio between mean activity evoked by contra- and ipsilateral stimuli:

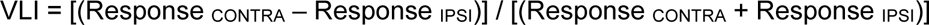

Positive values represent stronger responses to contralateral targets, and negative, to ipsilateral. We also determined with two sample, two-tail t-tests whether the distributions of responses for each neuron over trials was significantly different for contra-versus ipsilateral stimuli. We observed a broad range of laterality with 48 neurons (24% of visually facilitated neurons) producing significantly stronger responses to contralateral stimuli, 73 neurons (36% of visually facilitated neurons) producing significantly stronger responses to ipsilateral stimuli, and the remainder producing indistinguishable responses (**Fig. 3C**). In this sample, the distribution of VLI was modestly, but significantly less than 0 (mean VLI = −0.0505, t (202) = −2.404, p = 0.017). The average activity of neurons responding best to contralateral stimuli was weaker are arose later than the average activity of neurons responding best to ipsilateral stimuli (**Fig. 3D**).

The latencies of the facilitated responses ranged from 31 to 149 ms with mean ± SEM of 92.4 ± 1.9 ms (**Fig. 4**). One half of these cells exhibited latencies under 90 ms and by 112 ms, 75% of neurons were active. Significant variation in latency compared to previously reported areas was confirmed by a Kruskal-Wallis one-way ANOVA on ranks (H (9,883) = 400.31, p < 0.001). Multiple post-hoc Mann-Whitney two-way rank sum comparisons with Bonferroni correction were conducted to compare latencies between areas. Bonferroni corrected p-values reported henceforth are obtained by multiplying the unaltered p-value by the number of comparisons made. Combining neurons visually facilitated neurons recorded from both monkeys, this distribution (median = 90 ms) is indistinguishable from previously reported values in SEF (median = 76 ms; U(64,203) = 6981, p = 0.1395), and cingulate cortex (median = 94 ms; U(23, 203) = 2564, p = 1.0) (Pouget et al., 2005).

**Figure 4.**
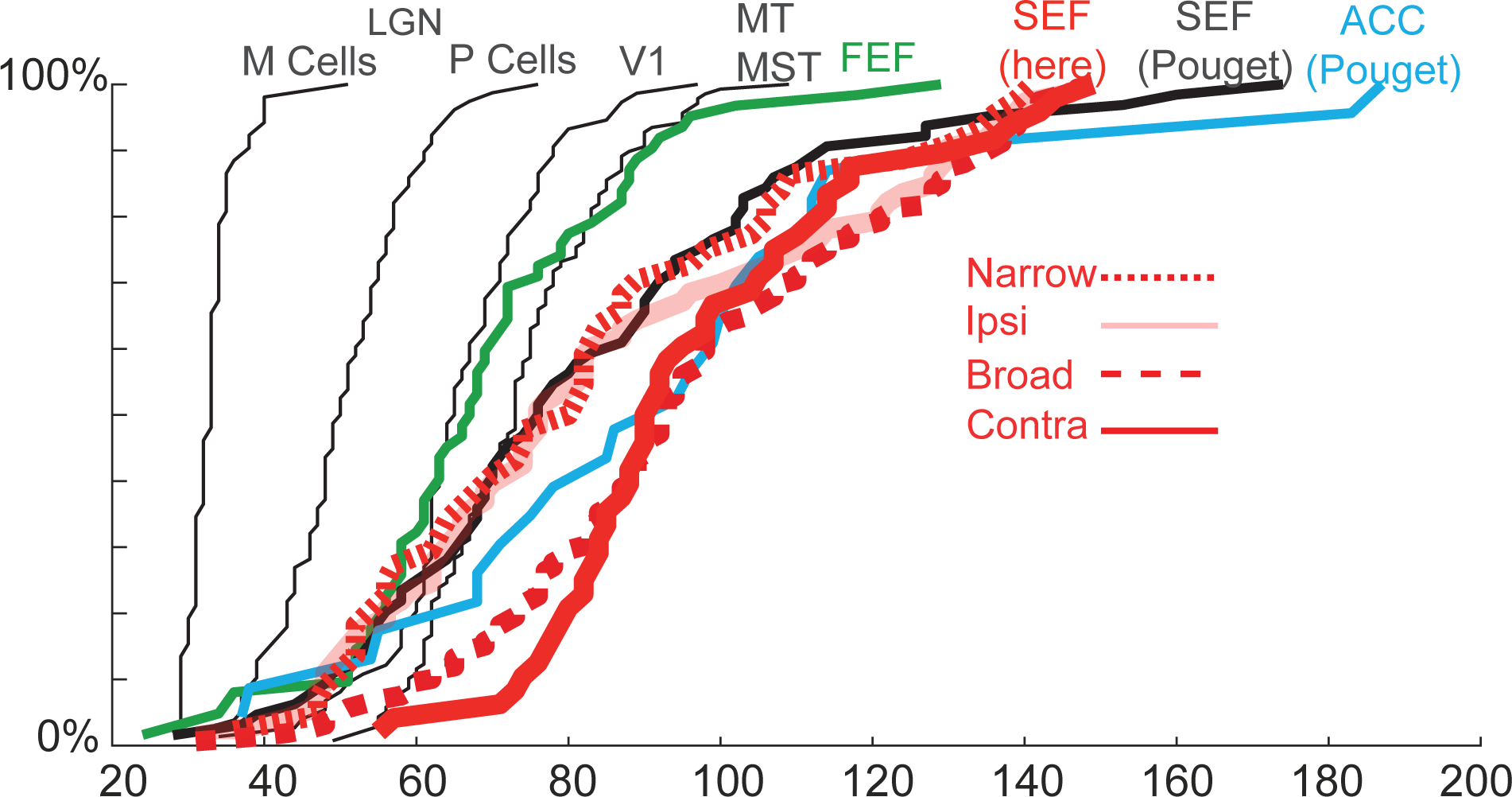
Cumulative distributions of visual responses sampled previously across the visual pathway (Schmolesky et al. 1998), of visual responses sampled previously in SEF (Pouget et al. 2005; Godlove et al. 2014), and of visual responses reported here.

The latencies of visual responses in SEF were significantly different than those measured in the lateral geniculate nucleus magno-(median = 33 ms; U(52, 203) = 1438, p < 0.001) and parvo-cellular (median = 50 ms; U(78, 203) = 4181, p < 0.001) layers, V1 (median = 65 ms; U(74, 203) = 5684, p < 0.001), MT and MST (median = 73 ms; U(138, 203) =1722, p < 0.001) and FEF (median = 73 ms; U(62, 203) = 4989, p < 0.001) (Schmolesky et al., 1998).

We observed idiosyncratic differences between monkeys with visual responses significantly faster in monkey Eu (median = 88ms; 88.5 ± 2.3 ms), compared to monkey X (median = 98 ms; 101.6 ± 3.4 ms; U(143,60) = 13457, p = 0.0031).

Neurons with broad spikes responded after a mean ± SEM latency of 95.4 ± 2.1 ms, while neurons with narrow spikes responded after 81.6 ± 4.2 ms. The visual response latency of neurons with broad spikes (median = 93.0 ms) was significantly different than that of neurons with narrow spikes (median = 80.5 ms; U(159,44) = 17293, p = 0.0018).

Neurons with stronger contralateral responses responded with a median latency of 90 ms, while neurons with stronger ipsilateral responses responded with a median latency of 75 ms. Neurons which did not display a laterality preference had a median latency of 94ms. The visual response latency of neurons with stronger responses for contralateral targets was significantly different than that of neurons with stronger responses for ipsilateral targets (U(48,73) = 3459, p = 0.005).

### Functional architecture of visual responses in supplementary eye field

In 16 recording sessions, penetrations were perpendicular to the cortical layers, verified through combined MR and CT imaging (Godlove et al., 2014). In these penetrations, we sampled 293 neurons, of which 119 (41%) produced facilitated responses. Visually-responsive neurons were observed across all cortical layers (**Fig. 5**).

**Figure 5.**
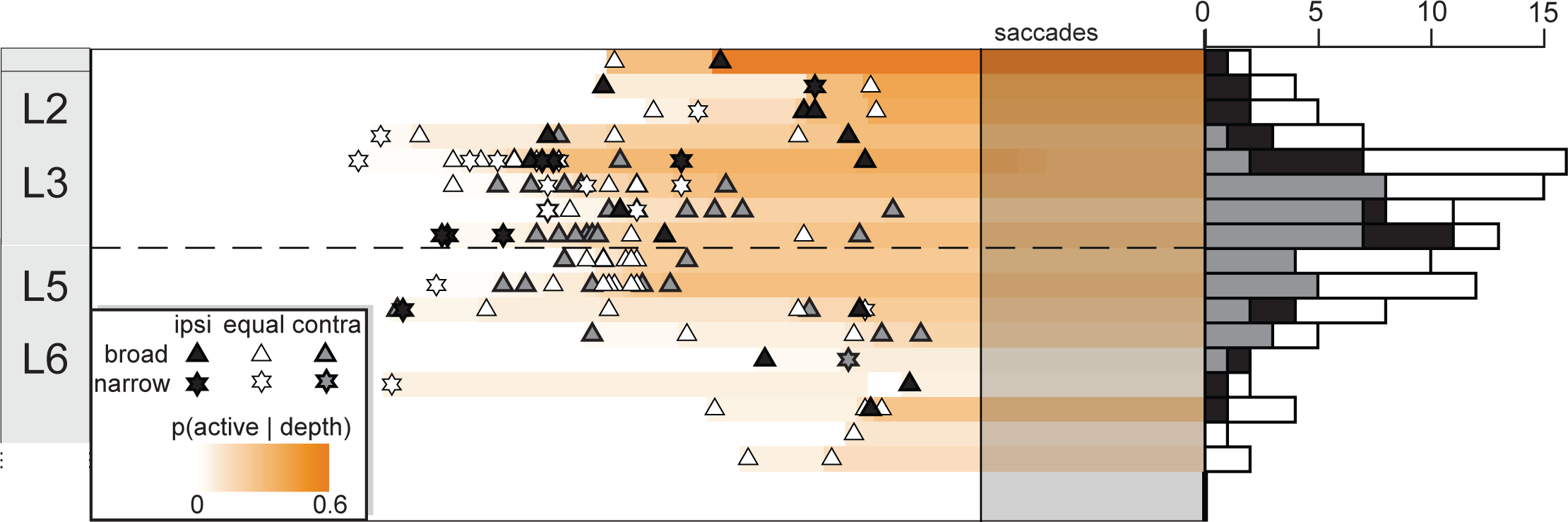
A. Time-depth plot showing latency and recruitment of facilitated visual responses across depth. Symbols mark beginning of visual response for neurons with spike width ≥ 250 µm (triangles) and neurons with spike width < 250 µm (stars). Color map indicates the percentage of neurons with visual responsiveness through time at each depth relative to the sampling distribution. Dashed horizontal line marks L3–L5 boundary. The lower boundary of L6 is not discrete. B. Laminar distribution of visual response laterality. Contralateral-preferring neurons have an LSI > 0 whereas ipsilateral-preferring neurons have an LSI < 0.

Time-depth plots of the recruitment of these neurons show that they are encountered most in layers 2 and 3 and less frequently in layer 5 and 6. Based on the distribution of neurons sampled across the layers, visual neurons were observed at a less-than-expected incidence in layer 5 and 6 (χ^2^ = 12.599, p = 0.006). The latency of visual responses was different between cortical layers using a one-way ANOVA (F(3,115) = 7.109, p < 0.001). Differences between pairs of cortical layers were determined using a pairwise t-test with Bonferroni correction. For pairwise comparisons, layers 2 and 3 (p = 0.040), layers 3 and 6 (p < 0.001) and layers 5 and 6 (p = 0.027) had significantly different onset times. Consistent with the canonical cortical microcircuit, middle layers 3 (90.7 ± 2.6 ms) and 5 (100.0 ± 3.2 ms) were activated first before layers 2 (107.9 ± 3.7 ms) and 6 (124.7 ± 3.9 ms). However, like Godlove et al. (2014), our findings were inconsistent with the canonical cortical microcircuit, as layers 3 and 5 appear to be coactivated (p > 0.99). As described above, the visually-evoked activity persisted until after the saccade with a significant difference in termination time across layers (F(3, 115) = 4.775, p = 0.004). Layers 2 and layer 5 had significantly different termination times (p = 0.002).

Neurons with stronger responses to ipsilateral targets were distributed through the cortical layers differently than those with stronger responses to contralateral targets using a chi-squared test, excluding neurons without a laterality preference (χ^2^ = 16.42, p <0.001). Multiple post-hoc pairwise chi-square tests with Bonferroni correction were conducted. Although observed in similar proportions in L2 and L6 (χ^2^ = 0.2778, p = 1.0) neurons with stronger ipsilateral responses were more prevalent in L2 (L2 vs. L3 χ^2^ = 9.070, p = 0.0026; L2 vs. L5 χ^2^ = 11.74, p = 0.0037) (**Fig 5**).

We calculated the visual lateralization index for neurons in each layer. Whereas L2 (mean LSI = - 0.0834) and L6 (mean LSI= −0.0850) had on average negative LSI, L3 (mean LSI = 0.1070) and L5 (mean LSI = 0.1072) had a positive LSI. Neurons in L2 preferred ipsilateral targets, and neurons in L5 preferred contralateral targets (F (3, 115) = 4.3972, p = 0.0057). Furthermore, whilst nearly all neurons with stronger contralateral responses had broad spikes (45/48 broad, 94%), neurons with stronger ipsilateral responses were a mix of broad (56/73, 63%) and narrow spikes.

To characterize further the laminar organization of visual sensitivity in SEF, we calculated current source density (CSD) from the recordings of local field potentials in response to contra- and ipsilateral targets (**Fig 6**). Several observations merit consideration. First, in contrast to the transient CSD evoked by light flashes in a passive state (Colonius et al., 2001), the CSD evoked by the target during the countermanding saccade task were prolonged until after saccade initiation. Second, the magnitude of the sinks produced by target presentation during the task were notably greater than those produced by flashes of light during a passive state with no reward (Godlove et al., 2014).

**Figure 6.**
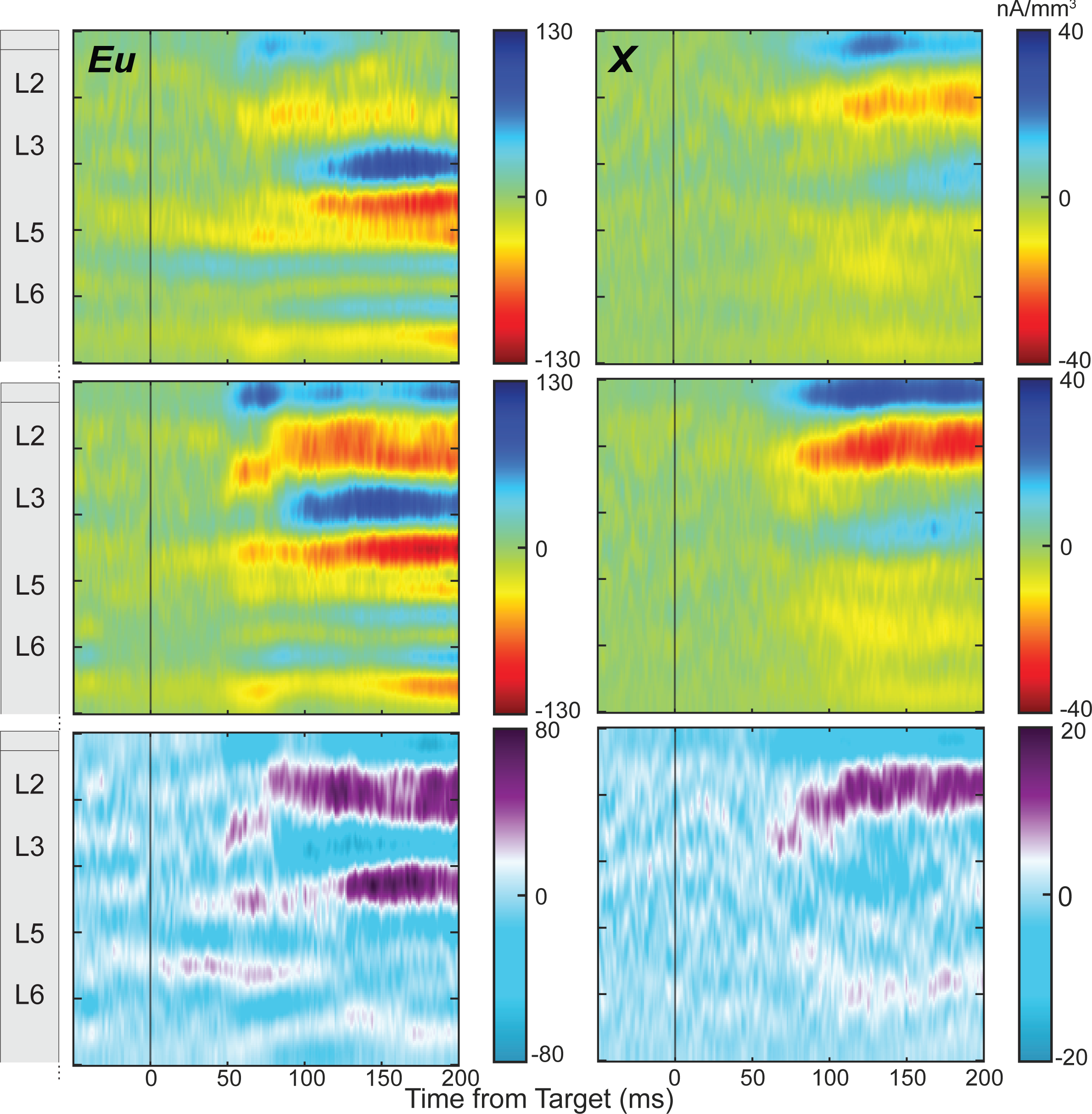
Visually evoked current-source density for contralateral target (top), ipsilateral target (middle), and their difference (bottom).

Third, a characteristic pattern of current sinks was observed across monkeys, with an early sink in L3 after ∼50-75 ms followed by a prolonged sink in upper L3 and L2 sustained until saccade initiation. Comparing the CSD in the passive condition to that reported here, the first sink initiates in layer 3 ∼50 ms after the visual target appeared, followed almost immediately by a second sink in layer 5. The third sink in the passive visual condition occurred in layers 1 and 2 ∼150ms after visual target appearance. Due to the prolonged CSD in response to the visual target, the third sink observed in response to the passive flash stimulus merges with the first sink with noticeable peaks in the prolonged sink. Finally, a fourth sink in layer 6 was observed in both the countermanding task and during passive stimulation. In response to the visual target, this current sink coincided with the second sink but in response to the passive visual flash, this sink coincided with the later third sink. Fourth, differences were observed across monkeys. Relative to monkey X, the CSD observed in monkey Eu had more pronounced sinks in L3 and L6. Fifth, the current sinks were stronger for ipsilateral targets, particularly in layers 2 and 3.

## DISCUSSION

These results provide the first description of the laminar organization of task-related visual responsiveness in SEF, complementing our previous description of the laminar organization of responses to passive visual flashes (Godlove et al., 2014). These results complement our earlier descriptions of the laminar organization of neurons signaling errors and reward gains and losses (Sajad et al., 2019) and signaling conflict, event timing and goal maintenance (Sajad et al., 2022) and reinforce the basic differences between granular sensory and agranular cortical areas (Ninomiya et al., 2015). These data reveal an unexpectedly pronounced preference for ipsilateral visual stimuli that calls for further investigation.

### Visual responses in the supplementary eye field

Around 40% of neurons recorded from SEF responded to presentation of visual targets. Previous studies have reported anywhere from ∼10% to 70% of visually responsive neurons (Schlag and Schlag-Rey, 1987; Schall, 1991a, b; Russo and Bruce, 2000; Roesch and Olson, 2003; Nakamura et al., 2005; Pouget et al., 2005; Purcell et al., 2012; Godlove et al., 2014; Bharmauria et al., 2021), albeit with vastly different neuron inclusion criterion. Visually responsive neurons in SEF have been demonstrated to be either visually facilitated or suppressed (Schlag and Schlag-Rey, 1987). However, like Schall (1991a), we report few visually suppressed neurons. Differences across studies are likely due to differences in single-unit sampling and in strategies for characterizing neurons based on research goals. Based on this new information from unbiased sampling with a linear microelectrode, another source of differences across studies can be the depth of sampling. With the linear electrode array, visually responsive neurons were significantly more common in layers 2 and 3 relative to layers 5 and 6.

The latencies of these neurons are consistent with those reported in previous studies in SEF (Pouget et al., 2005; Godlove et al., 2014). The visual response latencies in SEF followed those in the LGN, caudal visual areas, and FEF (Schmolesky et al., 1998) but were indistinguishable from those measured in cingulate cortex (Pouget et al., 2005).

These findings are generally consistent with previous studies demonstrating in SEF, demonstrating visual responses of single neurons (Schlag and Schlag-Rey, 1987; Schall, 1991a, b; Russo and Bruce, 2000; Roesch and Olson, 2003; Nakamura et al., 2005; Pouget et al., 2005; Purcell et al., 2012; Godlove et al., 2014; Bharmauria et al., 2021), with a few conspicuous differences. Based on anatomical connectivity, the visual responses in SEF can be delivered in afferents from the frontal lobe (frontal eye field and adjacent prefrontal areas), the parietal lobe (lateral intraparietal area and area 7a), and the temporal lobe (medial superior temporal area and superior temporal polysensory area) (Andersen et al., 1990; Huerta and Kaas, 1990; Schall et al., 1993). Another possible source is afferents from the lateral sector of the mediodorsal nucleus of the thalamus (Huerta and Kaas, 1990; Shook et al., 1991). The hypothesis about visual afferents from the mediodorsal nucleus is motivated by the observation that this nucleus conveys visual signals to the frontal eye field (Sommer and Wurtz, 2004). However, the nature of the signals conveyed from the mediodorsal nucleus to SEF are not known. SEF is also innervated by other thalamic nuclei, including the ventral anterior nucleus, nucleus X, the posterior subdivision of the ventral lateral nucleus, the central lateral nucleus, the parafascicular nucleus, and the suprageniculate-limitans nucleus (Matelli et al., 1989; Huerta and Kaas, 1990; Shook et al., 1991), but the visual responsiveness of neurons in those nuclei is untested.

Previous studies of SEF visual responses have found that receptive fields are most common in the contralateral hemifield (Schall, 1991a, b; Russo and Bruce, 2000; Purcell et al., 2012), but the ipsilateral representation exceeds that in FEF (Schall, 1991a, b). Yet, the preponderance of stronger responses to ipsilateral stimuli in the upper layers has not been reported before. This preponderance in the most superficial layers may have been overlooked in older studies, such as Russo and Bruce (2000), with single electrodes because of the spreading depression caused by the entry of the electrode (see review, Kramer et al. (2016)) and the possibility of puncturing immediately to deeper layers. Sampling across all cortical layers with linear multi-electrode arrays, after sufficient time to allow stabilization of the tissue and abeyance of spreading depression, we observed a higher incidence of neurons with stronger responses to ipsilateral stimuli concentrated in the upper cortical layers. Ipsilateral visual hemifield preference might also have been overlooked due to differences in the paradigm used to elicit visual activity. For instance, Purcell et al. (2012) noted that the mean response of SEF neurons decreased during a visual search task relative to a change detection task (see also Nakamura et al. (2005) for compatibility task). Additionally, while they observed a few ipsilateral target responses during the change detection task, this response virtually disappeared during visual search. Roesch and Olson (2003) also noted a similar phenomenon, where changing the reward amount altered visual responsiveness as well as directionality preference. The presence of stronger ipsilateral responses in SEF is explained by the pronounced connections of SEF with the SEF and FEF in the other hemisphere (Huerta et al., 1987; Huerta and Kaas, 1990) plus inputs from neurons representing both the contra- and ipsilateral visual fields in parietal area 7a (Motter and Mountcastle, 1981) and the superior temporal polysensory area (Bruce et al., 1981). Additionally, the prefrontal cortex, which has extensive reciprocal connections with SEF (Huerta and Kaas, 1990) also responds to visual information in the ipsilateral hemifield (Wimmer et al., 2016). The finding of stronger ipsilateral preferences in L2/3 is consistent with the density of supragranular neurons labeled by tracer injections in the opposite FEF (Huerta and Kaas, 1990). At the same time, we noted a strong ipsilateral preference in layer 6 for monkey Eu that were absent in monkey X when comparing between ipsilateral and contralateral CSDs. Stronger CSDs for monkey Eu than for monkey X was also observed in Godlove et al. (2014). Differences in laminar profiles of monkeys can be due to sex and species differences as monkey Eu was a male bonnet macaque (*Macaca radiata)* and monkey X was a female rhesus macaque (*Macaca mulata)*. Heterogeneity across SEF is another possible explanation based on differences in the neuron types found in different cortical columns (Sajad et al., 2019; Sajad et al., 2022).

### Visual responses when passive and performing a task

These descriptions of visual responses during a rewarded task contrast usefully with earlier reports describing the laminar organization of visual activity in the supplementary eye field in response to passive visual flashes (Godlove et al., 2014). First, this earlier study reported an equal portion of visually evoked neurons with spike facilitation and suppression. However, here we report the vast majority (90%) of visually responsive neurons exhibited enhancement to a visual target, whilst very few (10%) demonstrated significant suppression. Second, response latencies between cortical layers do not significantly differ during passive viewing. However, in the present study we observed visual neurons were first active in middle layers 3 and 5 followed by neurons in layers 2 and 6. Furthermore, although (Godlove et al., 2014) reported no difference in latency of putative pyramidal neurons and interneurons, we observed significantly shorter latencies in neurons with narrow relative to broad spikes. Finally, whilst the time-depth pattern of spiking activity and CSD were incongruent during passive viewing, during the active task both spiking and CSD activity during the countermanding task were both first active in layer 3 and 5.

Given that data from the passive visual evoked potential task and countermanding task were recorded from the same monkeys within the same session, the differences in neural activity in the SEF between passive flashes and the countermanding task must be due to differences in the cognitive demands and engagement of the two conditions. In particular, during the passive task, gaze was not constrained and behavior, unrelated to reward. In contrast, during the countermanding task, gaze was necessarily controlled with strong contingency on reward.

## Conclusion

This description of the laminar structure of visual responses during a countermanding saccade task offers new information about the functional architecture of SEF. The findings contrast with an earlier description of the passive visual responses in SEF (Godlove et al., 2014) and further confirm basic differences between this exemplar agranular cortical area and the canonical cortical microcircuit derived from primary visual cortex (Ninomiya et al., 2015). The findings complement our previous reports of the laminar organization of neurons involved with performance and reward monitoring (Sajad et al., 2019) as well as conflict, event timing, and goal maintenance (Sajad et al., 2022).

